## Supplemental Table 1 for "Structural insights into loss of function of a pore forming toxin and its role in pneumococcal adaptation to an intracellular lifestyle"

**S1 Table. Diffraction data collection and structure refinement statistics**

| **DATA COLLECTION STATISTICS** ^A^ | **PLY-NH** |
| --- | --- |
| Wavelength(Å) | 0.95372 |
| Crystal to detector distance (mm) | 213.0 |
| Space group | *P*2_1_2_1_2_1_ |
| Unit cell parameters  *a*, *b*, *c*(Å)  α, β, γ (°) | 24.5, 84.7, 214.6  α = β = γ = 90 |
| Mosaicity(°) | 0.2 |
| Resolution limit(Å) | 38.3-2.2 (2.25-2.2) |
| Total No.of reflections | 280245 (89745) |
| Unique reflections | 23881 (7414) |
| Redundancy | 11.7 (12.1) |
| I/σ˂I˃ | 10.56 (2.7) |
| Completeness (%) | 99.9 (100) |
| R_merge_ (%)^b^ | 24.0 (131.8) |
| CC_1/2_ | 99.6 (84.3) |
| Overall B factor from Wilson plot (Å^2^) | 34.0 |
| **Refinement statistics** | |
| Working set | 22691 |
| Test set | 1195 |
| R_work_ (%)^c^ | 22.3 |
| R_free_ (%)^d^ | 26.3 |
| Total no. of residues | 469 |
| Total no. of water molecules | 273 |
| Overall B factor (Å^2^) | 38.05 |
| R.M.S.D. of bond length (Å) | 0.006 |
| R.M.S.D. of bond angle (°) | 1.156 |
| *Ramachandran plot analysis* | |
| Most favoured (%) | 95 |
| Additionally allowed (%) | 5 |
| Disallowed region (%) | 0 |
| PDB code | **6JMP** |

*^a^* Values in parentheses are for the highest-resolution shell

*^b^*
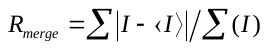
 I where is the observed integrated intensity, 〈*I*〉 is the average integrated intensity obtained from multiple measurements, and the summation is over all observed reflections.

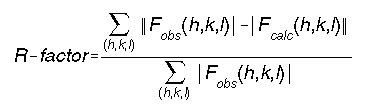
^c^ where F_obs_ and F_calc_ are observed and calculated structure factors respectively.

^d^R_free_ was calculated as for R_work_ but only 5% data left out of refinement procedure has been used in the calculation
