## Supplemental Table 1 for "Structural insights into loss of function of a pore forming toxin and its role in pneumococcal adaptation to an intracellular lifestyle"

**S2 Table. Relative hemolytic activities of cysteine substituted mutants of Ply-H and Ply-NH, before and after labelling with NBD dye.**

| **Conc. (µM)** | **0.1** | | **0.01** | |
| --- | --- | --- | --- | --- |
|  | **Unlabelled** | **NBD-labelled** | **Unlabelled** | **NBD-labelled** |
| **Ply-H mutants** |  | |  | |
| **S167C** | 110.3 | 107.0 | 108. 7 | 105.8 |
| **H184C** | 104.6 | 105.7 | 102. 4 | 107.1 |
| **D257C** | 108.6 | 109.7 | 105.5 | 102.7 |
| **E260C** | 109.4 | 101.8 | 100.9 | 100. 8 |
| **Ply-NH mutants** |  | |  | |
| **S175C** | 0.1 | 0.4 | 0.1 | 0.2 |
| **H184C** | 0.0 | 0.2 | 0.1 | 0.2 |
| **D257C** | 0.4 | 0.5 | 0.1 | 0. 3 |
| **E260C** | 0.5 | 0.1 | 0.4 | 0.1 |

Values shown are % hemolysis (with 100% representing complete lysis as obtained with Triton X-100).
