## Supplemental Table 3 for "Structural insights into loss of function of a pore forming toxin and its role in pneumococcal adaptation to an intracellular lifestyle"

**S3 Table. List of *S. pneumoniae* strains.**

| **STRAIN** | **SOURCE** |
| --- | --- |
| D39 (Serotype 2, encapsulated) | Mitchell TJ, Univ. of Birmingham, UK |
| R6 (Serotype 2, non-encapsulated) | Mitchell TJ, Univ. of Birmingham, UK |
| TIGR4 (Serotype 4, encapsulated) | Mitchell TJ, Univ. of Birmingham, UK |
| 01-1956 (Serotype 1, ST306, encapsulated) | Mitchell TJ, Univ. of Birmingham, UK |
| D39:Ply-H | This study |
| D39:Ply-NH | This study |
| D39Δ*ply* | This study |
| D39:Ply^W433F^ | This study |
| R6:Ply-H | This study |
| R6:Ply-NH | This study |
| R6:Ply-DM (Ply-NH^H150Y+I172T^) | This study |
| R6Δ*ply* | This study |
| R6:Ply^W433F^ | Surve *et al* (27) |
| R6:Ply-H:HlpA-GFP/tagRFP | This study |
| R6:Ply-NH:HlpA-GFP/tagRFP | This study |
