## Supplemental Table 4 for "Structural insights into loss of function of a pore forming toxin and its role in pneumococcal adaptation to an intracellular lifestyle"

**S4 Table. List of primers.**

| **NAME** | **TYPE** | **PRIMER SEQUENCE** |
| --- | --- | --- |
| E286A | Forward | 5′ CAATACAGCAGTGAAGGCGGTTATTTTAG 3′ |
|  | Reverse | 5′ CTTCACTGCTGTATTGTCCAAAATCTGC 3′ |
| V287A | Forward | 5′ CAATACAGAAGGGAAGGCGGTTATTTTAGG 3′ |
|  | Reverse | 5′ CGCCTTCCCTTCTGTATTGTCCAAAATC 3′ |
| K288A | Forward | 5′ CAGAAGTGGCGGCGGTTATTTTAGGGG 3′ |
|  | Reverse | 5′AACCGCCGCCACTTCTGTATTGTCCAAAATC 3′ |
| C428A | Forward | 5′ AGAGAGGCTACCGGGCTTGCCTGGGAATG 3′ |
|  | Reverse | 5′ CAAGCCCGGTCGACTCTCTAATTTTGACAG 3′ |
| H150Y | Forward | 5′ CCCAGCTAGAATGCAGTATGAAAAAATC 3′ |
|  | Reverse | 5′ GTGAGCCGTGATTTTTTCATACTGCATTC 3′ |
| I172T | Forward | 5′ GTTTGGTTCTGACTTTGAAAAGACAGGGAATTC 3′ |
|  | Reverse | 5′ GTTAAAATCAATATCAAGAGAATTCCCTGTCTTTTC 3′ |
| S167C | Forward | 5′ CTCAAGGTCAAGTTTGGTTGCGACTTTGAAAAG 3′ |
|  | Reverse | 5′ CCTGTCTTTTCAAAGTCGCAACCAAACTTG 3′ |
| S175C | Forward | 5′ GAAAAGATAGGGAATTGTCTTGATATTG 3′ |
|  | Reverse | 5′ GTTAAAATCAATATCAAGACAATTCCCTATC3′ |
| H184C | Forward | 5′ GATATTGATTTTAACTCTGTCTGTTCAGGCG 3′ |
|  | Reverse | 5′CTGCTTTTCGCCTGAACAGACAGAGTTAAAATC 3′ |
| D257C | Forward | 5′ GAAACCACGAGTAAGAGTTGTGAAGTAGAGGC 3′ |
|  | Reverse | 5′ CAAAAGCAGCCTCTACTTCACAACTCTTAC 3′ |
| E260C | Forward | 5′ GTAAGAGTGATGAAGTATGTGCTGCTTTTG 3′ |
|  | Reverse | 5′ CAAAGATTCAAAAGCAGCACATACTTCATC 3′ |
| *ply*-upstream | Forward | 5’ TATATAGGTACCGGAAATGTCTCAATCCAGCTA 3′ |
|  | Reverse | 5′ TATATACTCGAGCTTCTACCTCCTAATAAGTTCC 3′ |
| *ply*-downstream | Forward | 5′ TATATAGGATCCGAGAGGAGAATGCTTGCGAC 3′ |
|  | Reverse | 5′ AGTGGATCTAGACAGTTCTTATAGGCGCTATTGT 3′ |
| *ply* ORFs | Forward | 5′ CATCCTCTCGAGATGGCAAATAAAGCAGTA 3′ |
|  | Reverse | 5′ CTTATCGGATCCCTAATCATTTTCTACCTTATCC 3′ |
| Spectinomycin resistance cassette | Forward | 5′ ATCCGGATCCAATCTGATTACCAATTAGAATG 3′ |
|  | Reverse | 5′ CCGCGGATCCCATATATAATCTAGAATAAAATTAAC 3′ |
