## Supplementary figures and images for "Structural insights into loss of function of a pore forming toxin and its role in pneumococcal adaptation to an intracellular lifestyle"

### Supplemental Figure 1

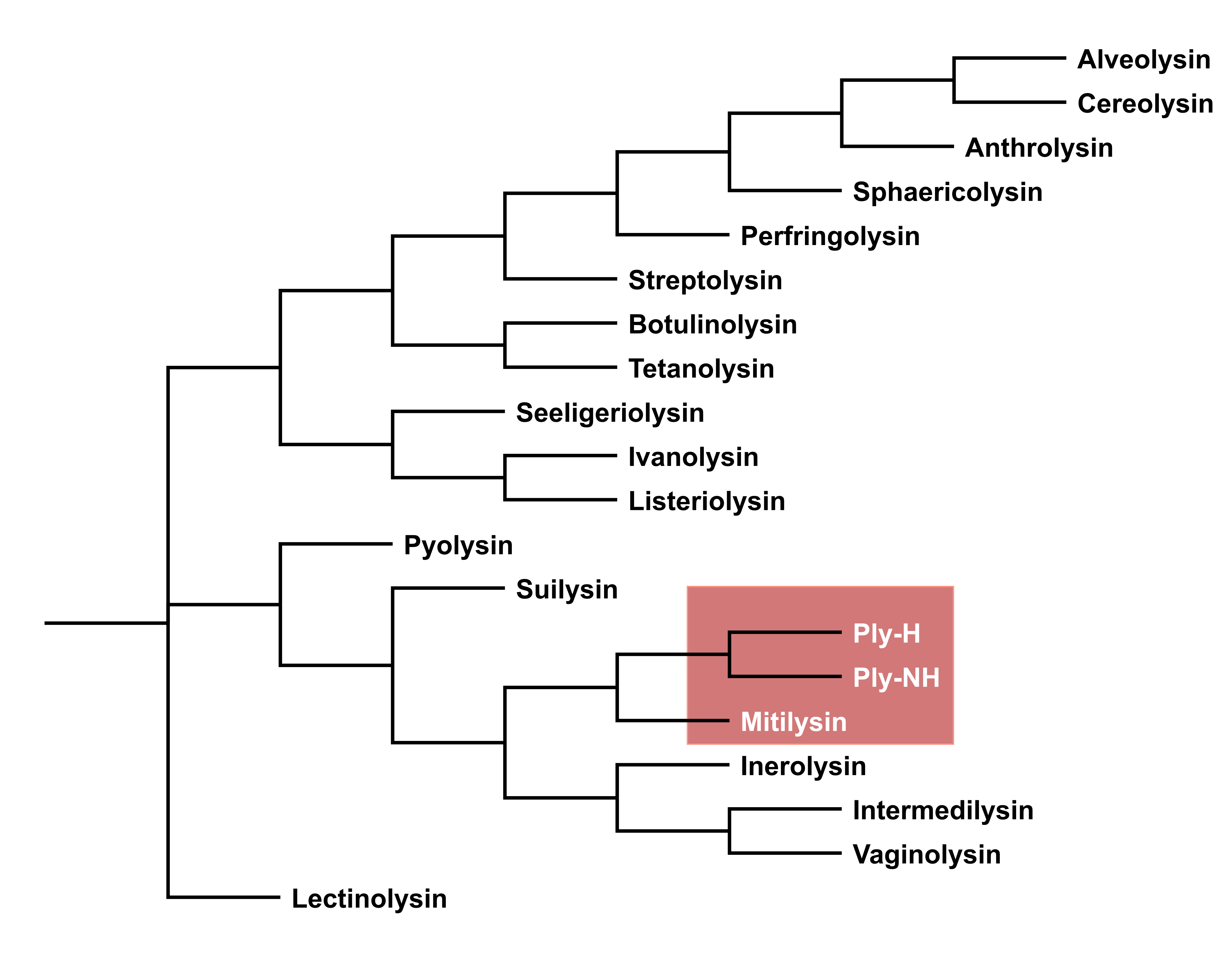

### Supplemental Figure 2

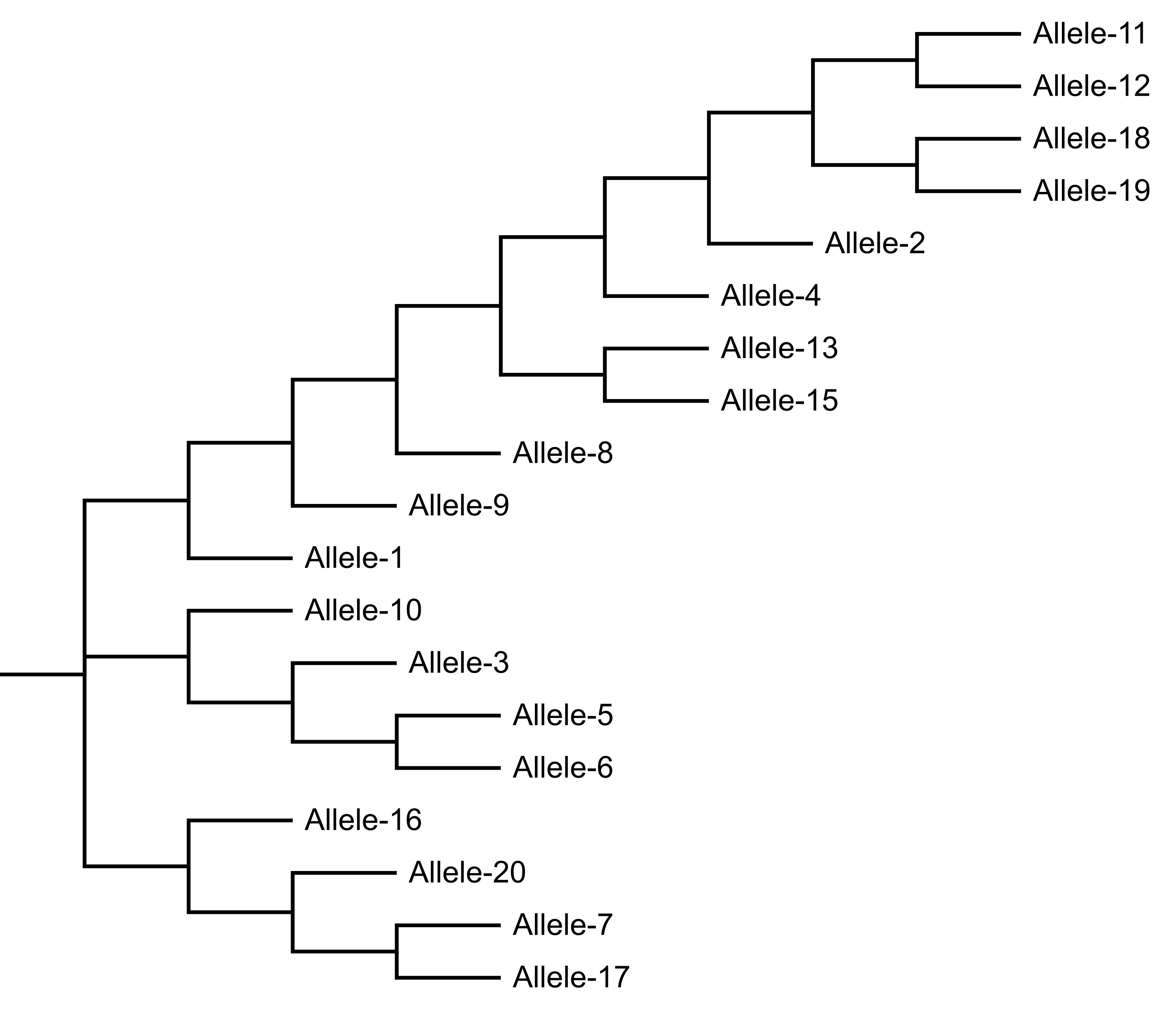

### Supplemental Figure 3

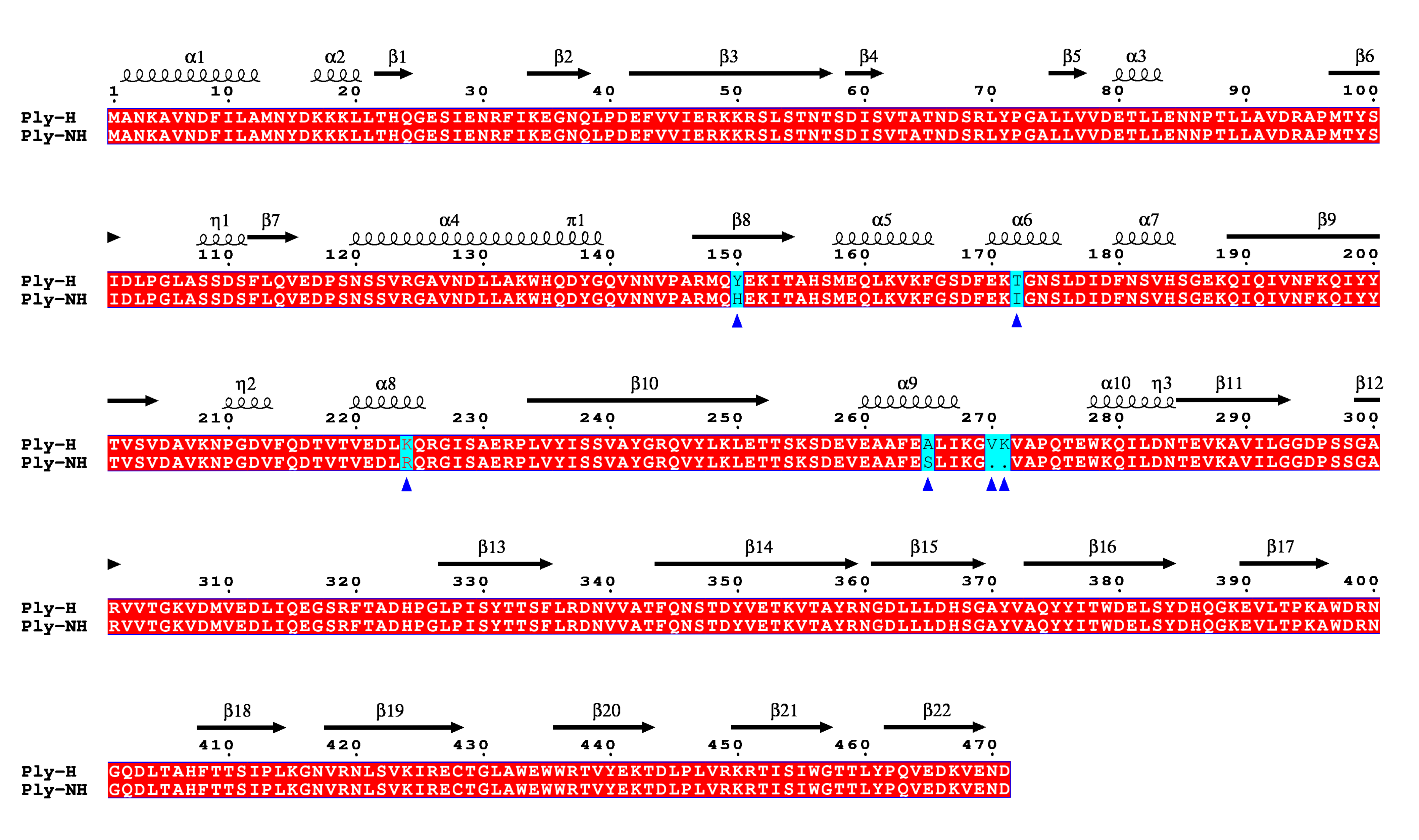

### Supplemental Figure 4

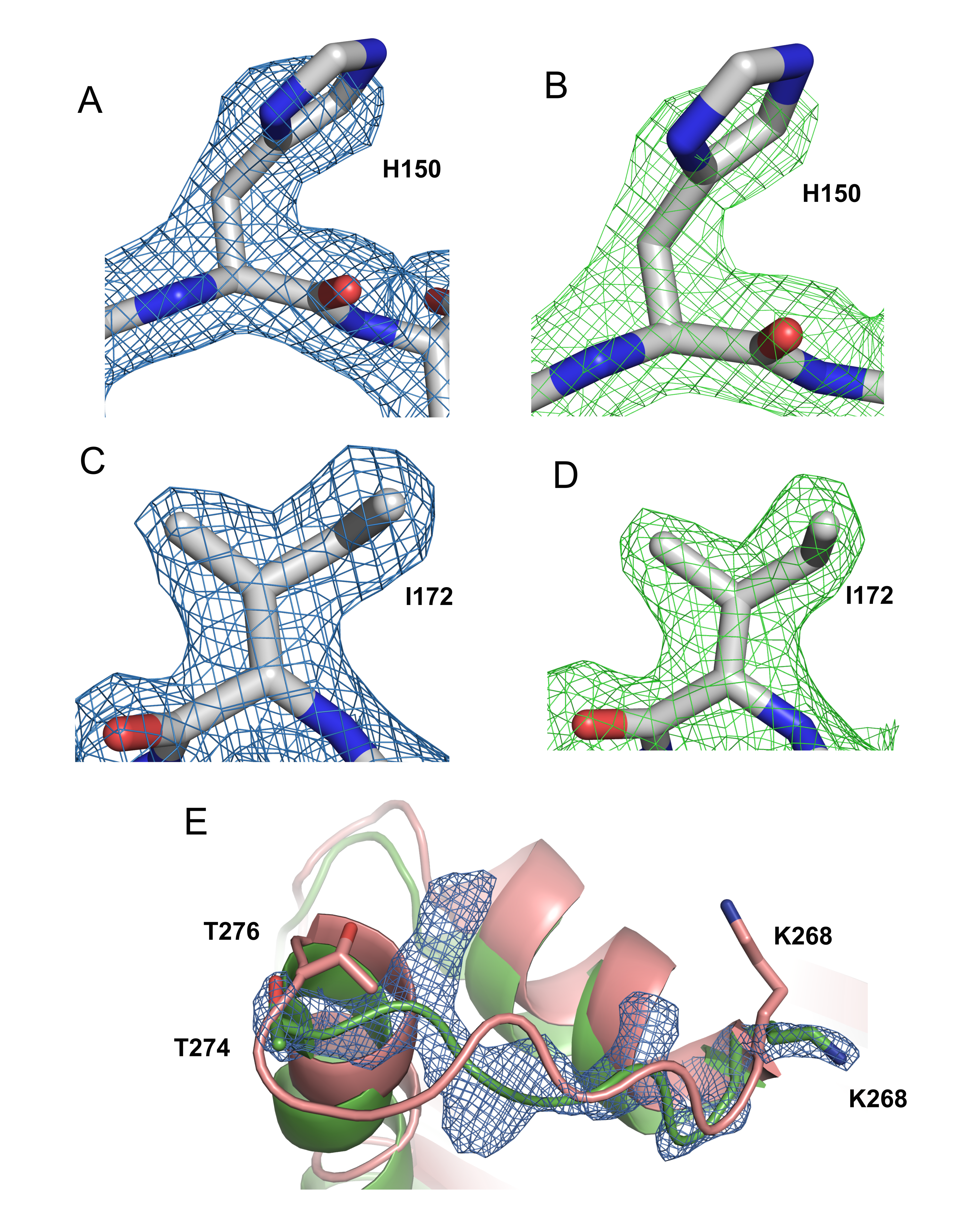

### Supplemental Figure 5

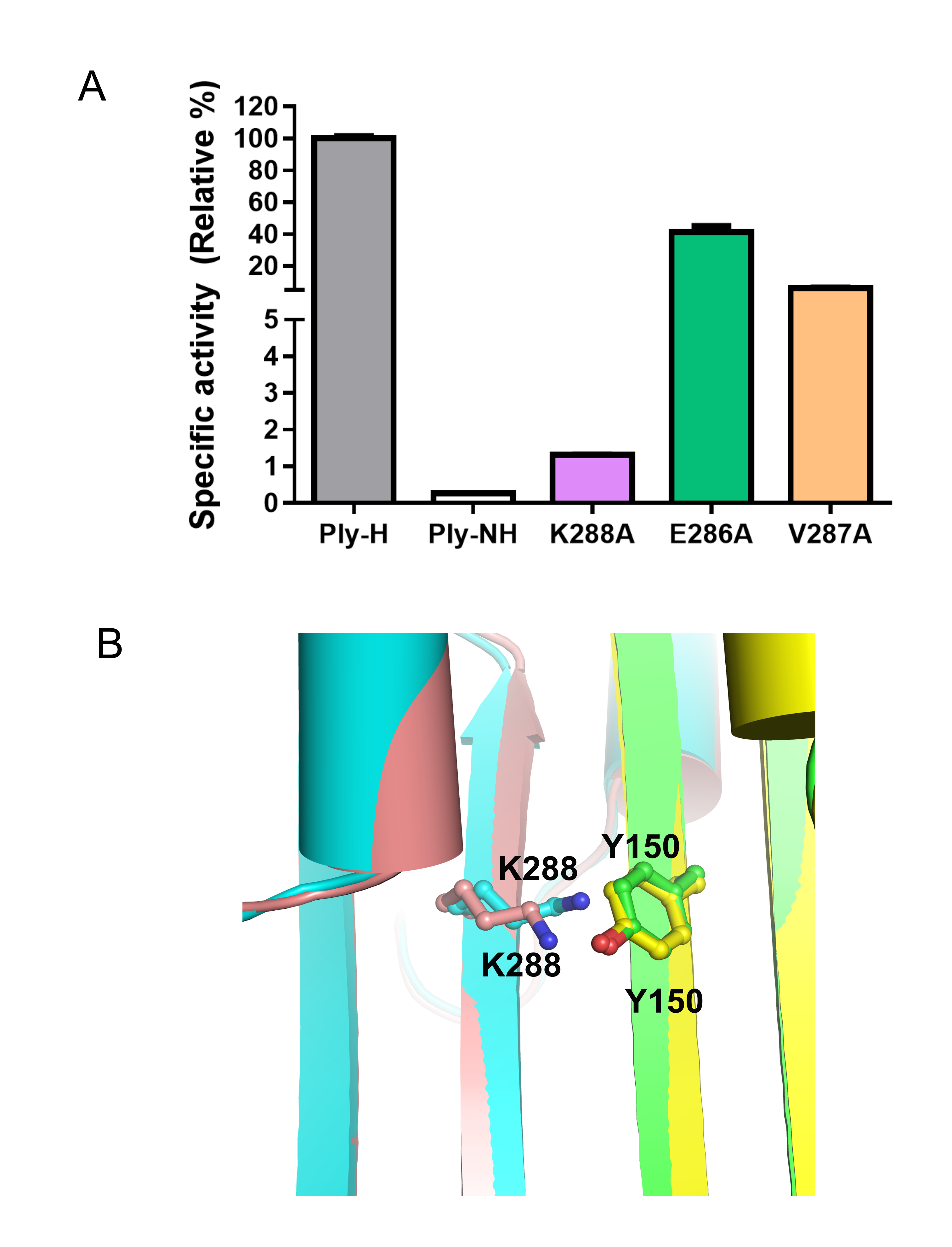

### Supplemental Figure 6

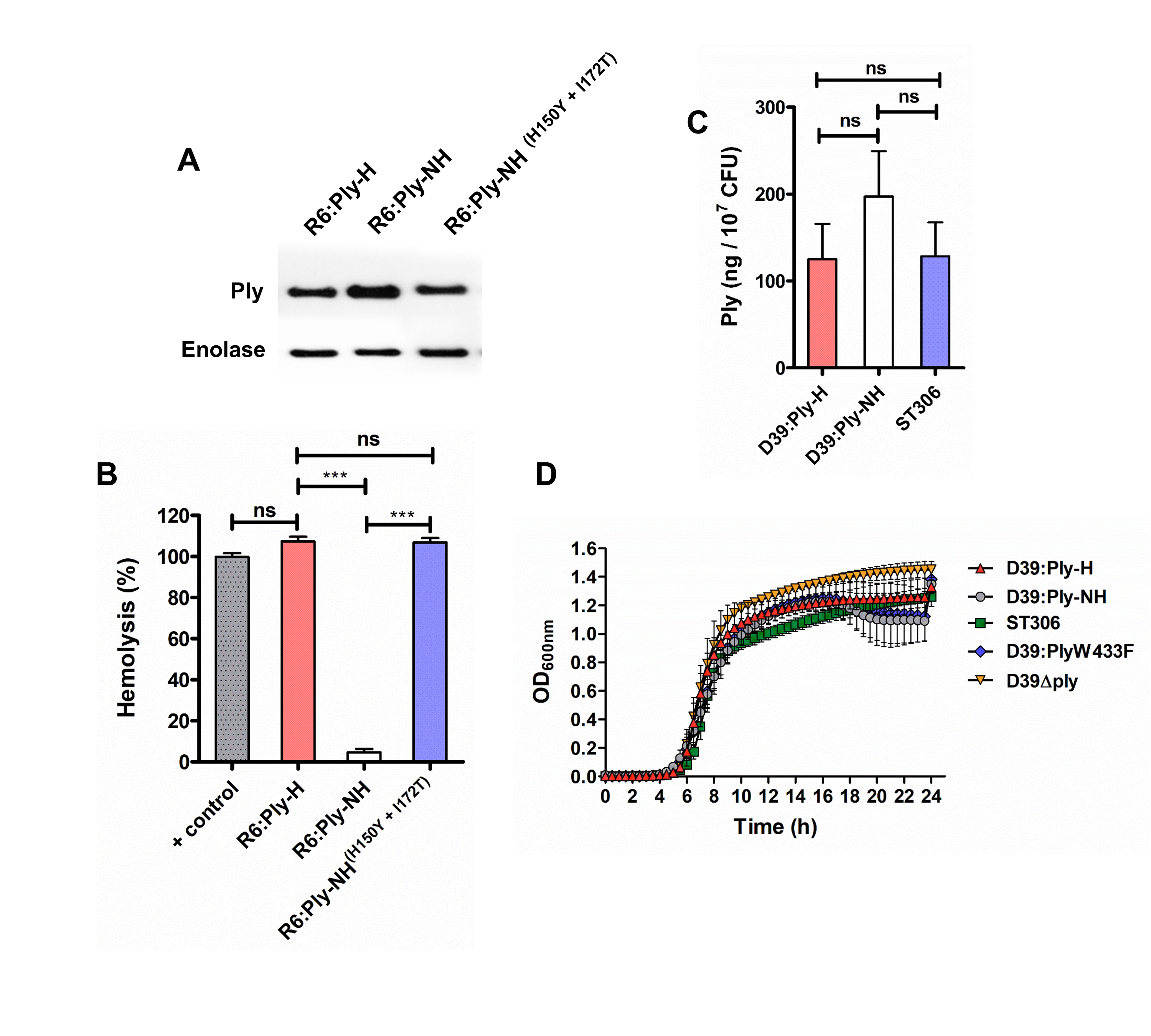

### Supplemental Figure 7

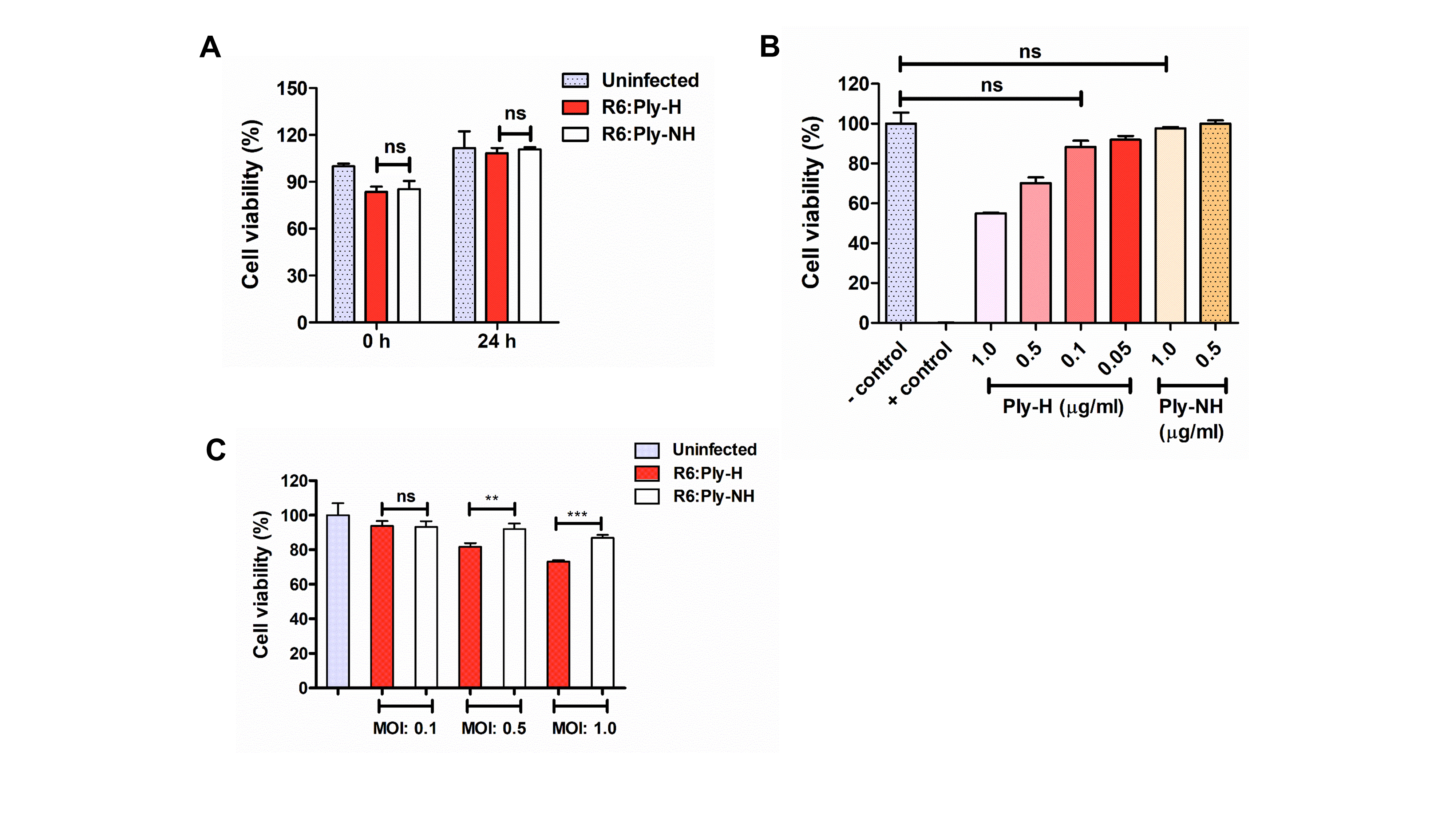

### Supplemental Figure 8

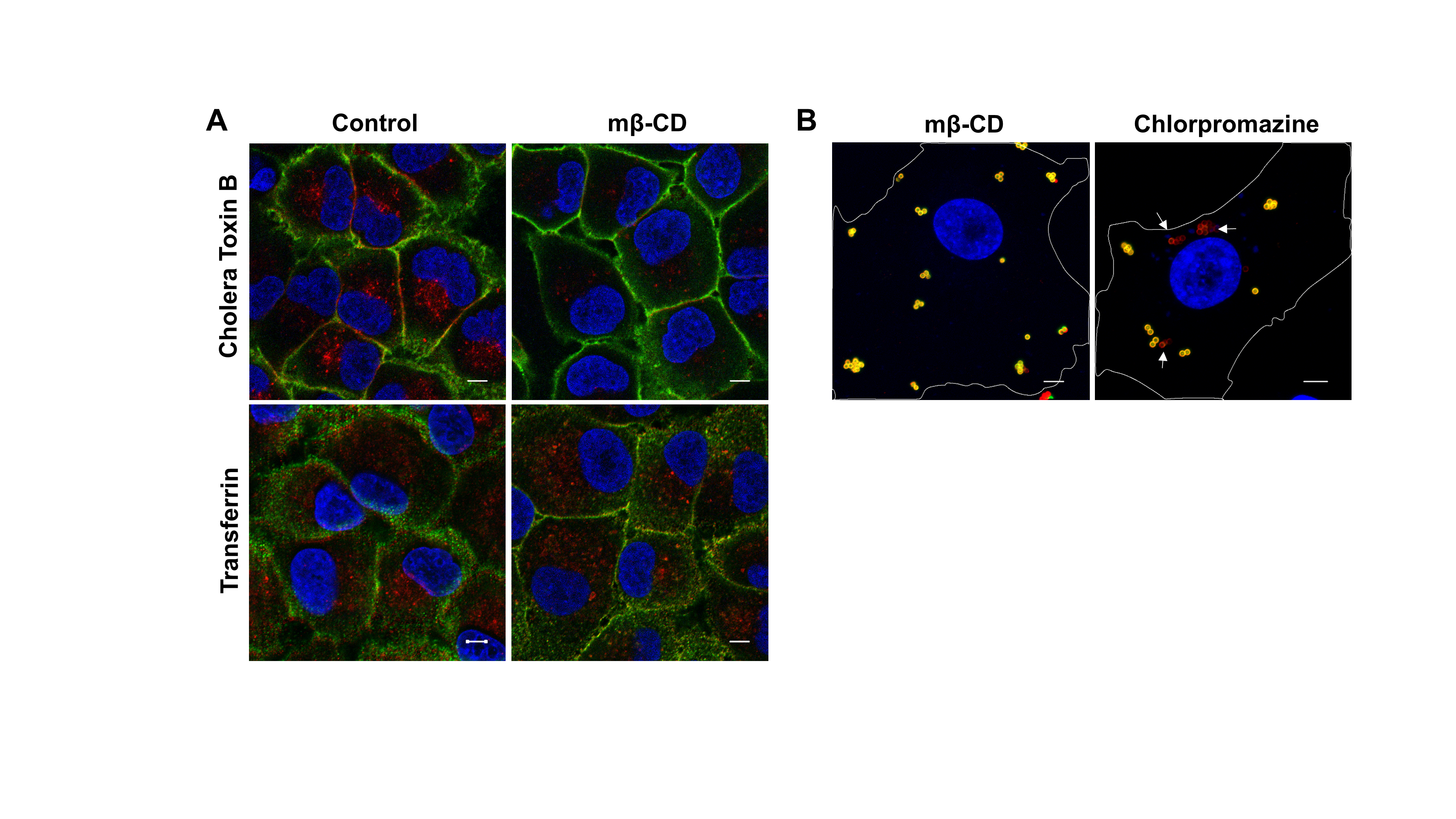

### Supplemental Figure 9

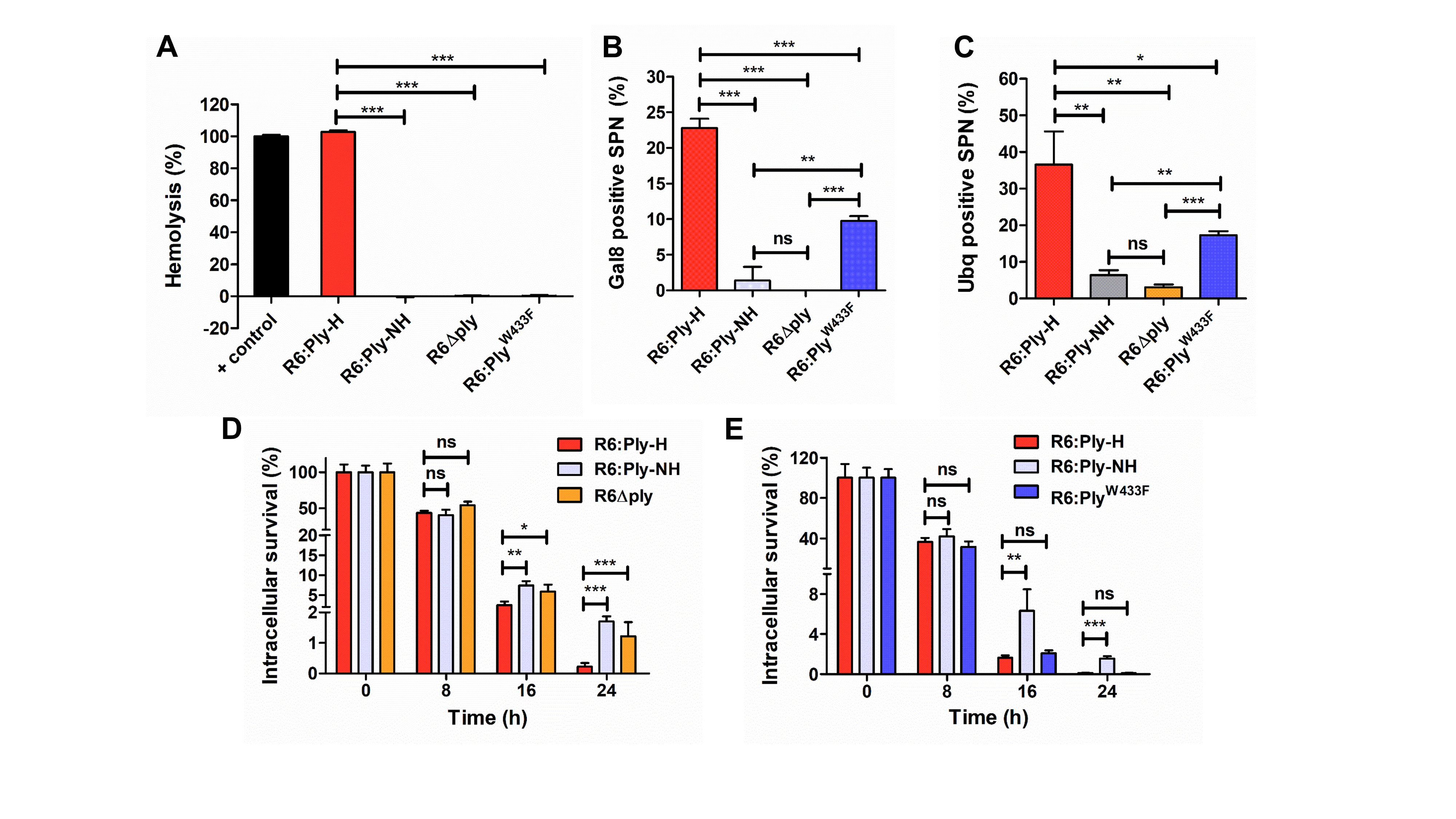

### Supplemental Figure 10

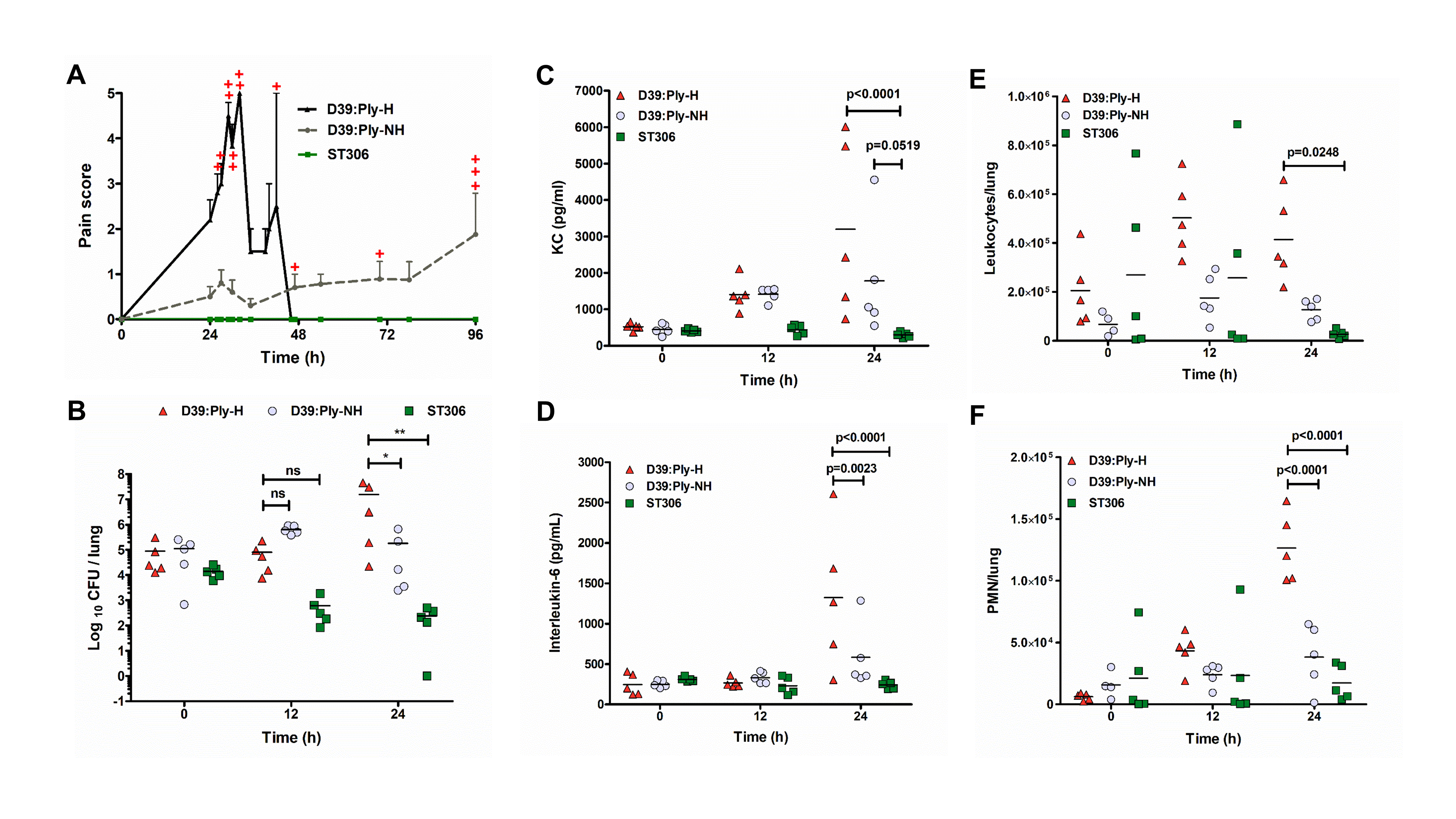
